# Epigenetic Variation in Tree Evolution: a case study in black poplar (*Populus nigra*)

**DOI:** 10.1101/2023.07.16.549253

**Authors:** Mamadou Dia Sow, Odile Rogier, Isabelle Lesur, Christian Daviaud, Emile Mardoc, Edmond Sanou, Ludovic Duvaux, Peter Civan, Alain Delaunay, Marie-Claude Lesage- Descauses, Vanina Benoit, Isabelle Le-Jan, Corinne Buret, Celine Besse, Harold Durufle, Régis Fichot, Grégoire Le-Provost, Erwan Guichoux, Christophe Boury, Abel Garnier, Abdeljalil Senhaji-Rachik, Véronique Jorge, Christophe Ambroise, Jorg Tost, Christophe Plomion, Vincent Segura, Stéphane Maury, Jérôme Salse

## Abstract

How perennial organisms adapt to environments is a key question in biology. To address this question, we investigated ten natural black poplar (*Populus nigra*) populations from Western Europe, a keystone forest tree of riparian ecosystems. We assessed the role of (epi)genetic regulation in driving tree species evolution and adaptation over several millions of years (macro-evolution) up to a few generations (micro-evolution). At the macro-evolution scale, polar experienced differential structural (gene loss) and regulation (expression and methylation) reprogramming between sister genomic compartments inherited from polyploidization events. More interestingly, at the micro-evolution scale, both genetic and epigenetic variations differentiate populations from different geographic origins, targeting specifically genes involved in disease resistance, immune response, hormonal and stress response that can be considered as key functions of local adaptation of long lifespan species. Moreover, genes involved in cambium formation, an important functional trait for forest trees, as well as basal functions for cell survival are constitutively expressed though methylation control. These results highlight DNA methylation as a marker of population differentiation, evolutionary adaptation to diverse ecological environments and ultimately opening the need to take epigenetic marks into account in breeding strategies, especially for woody plants.

## INTRODUCTION

Understanding the forces driving species evolution is central in biology. Evolution can be investigated at either short timescale, over a few hundreds of generations (so called micro-evolution) typically between different accessions of a species, or over long timescales of several millions of years between different species (macro-evolution). Genetic variations have long been considered as the major marker of these two facets of evolution, with specific approaches to study the extent of population dynamics and adaptive evolution at the micro-evolutionary scale (Messer et al. 2016) or lineage diversification and speciation from founder ancestral genomes at the macro-evolutionary scale (Murat et al. 2012, Pont and Salse, 2017). New discoveries over the past two decades suggest that epigenetic variation, precisely DNA methylation, also play a role in micro- and macro-evolutionary changes (Jablonka 2017).

At a macro-evolutionary timescale, flowering plants diversified from an ancestor suggested to be made of 15 protochromosomes with 22,899 protogenes dating back to 214 million years ago during the late Triassic era (Murat et al. 2017). Polyploidization (also known as whole genome duplication, WGD) has been proposed as a key mechanism through which new genetic material is generated during evolution promoting morphological and phenotypic biodiversity and contributing to the evolutionary success of the modern angiosperm species (Jaillon et al. 2007; Schnable et al. 2011; Pont et al. 2013). In that context, regulation of duplicated genes though DNA methylation may therefore represent a central phenomenon in the acquisition of novel functions and ultimately phenotypes in the newly formed polyploids compared to their diploid progenitors (El Baidouri et al. 2018; Wang et al. 2017; Chen et al. 2015; Davis et al. 2015; Jablonka 2017).

At a micro-evolutionary timescale, natural selection contributed to shape heritable epigenetic (epialleles) and genetic (alleles) variations in natural populations (Furrow and Feldman, 2013, Li et al. 2014). For instance, in the model plant *Arabidopsis thaliana*, inheritance of epialleles has been associated with population structure and heritable phenotypic variation including adaptive traits (Cortijo et al. 2014). DNA methylation represents one of the most stable epigenetic marks, corresponding to an addition of a methyl group in 5’ of cytosines. In plants, methylation occurs in all cytosine contexts, *i.e.*, in CpG, CHG and CHH where H could represent A, T or C. Our understanding of the role of DNA methylation in plants has been mostly obtained from the model plant *Arabidopsis thaliana*, where DNA methylation in the coding regions of genes drives medium to high levels of expression and targets mainly housekeeping gene functions, while the methylation in the promoter region of genes is generally associated with gene silencing (Zhang et al. 2006; Maunakea et al. 2010; Bewick and Schmitz 2017). Despite a growing number of evidences unravelling the relationship between DNA methylation and gene expression, further efforts are needed to better understand methylation control over gene expression regarding different genomic contexts, and the stability of such epigenetic changes through time. Moreover, the contribution of DNA methylation in the evolution and adaptation of long lifespan species, such as trees, remains an open question (Amaral et al. 2020).

In contrast to annual plants, trees are long-living organisms that are exposed to environmental challenges over their entire lifespan (Allen et al. 2010; Anderegg et al. 2016). They have developed various stress sensory mechanisms and physiological strategies to cope with fluctuating environmental changes. The difference between annual (herbaceous) and perennial species in genome evolution is exemplified by the expansion of disease-resistance (R) genes in long-lived species as a key evolutionary process to face a wide range of abiotic and biotic threats over their lifespans (Noir et al. 2001; Ribas et al. 2011; Plomion et al. 2018; Khan and Korban, 2022). Moreover, because of relaxed purifying selection, R-genes in trees were found to display higher levels of genetic diversity (Plomion et al. 2018). It has also been shown that the rate of epigenetic mutation is much higher than genetic variations and then represents a source for short-term adaptation especially for long-living species (van der Graaf et al. 2015).

Poplar is considered as a model forest tree species. Poplars are characterized by a high genetic diversity, fast juvenile growth, vegetative propagation capacity, and the genome of several species has been sequenced (Tuskan et al. 2006; Mader et al. 2016). Over the past decade, poplar has been widely used to investigate the role of DNA methylation in phenotypic plasticity and adaptation to environmental changes (Brautigam et al. 2013; Lafon-Placette et al. 2013, 2018; Zhu et al. 2013; Plomion et al. 2016; Conde et al. 2017; Le Gac et al. 2018, 2019; Sow et al. 2018a, 2018b, 2021; Vigneaud et al. 2023). However, several open questions remain to be addressed, such as (i) does epigenetics play a role in genome reprogramming in response to polyploidization events? (ii) are populations epigenetically differentiated and to what extent does epigenetic differentiation may differ to genetic differentiation? (iii) Can local adaptation be tracked at the epigenetic level?

## RESULTS

### Genomic pattern of DNA methylation in black poplar

Poplar methylome on a genome-wide scale was investigated from whole genome bisulfite sequencing (WGBS) carried out on ten natural populations, covering the geographical distribution of *Populus nigra* in Western Europe (Figure 1a). DNA methylation in CpG context (averaging 32.1% to 35.4%) appeared more frequent compared to CHG (from 20.0% to 21.9%) and CHH (4.4% to 6.5%) contexts (Figure S1, Table S2). Methylated cytosines or hereafter called SMPs (for Single Methylation Polymorphisms, Figure 1b) were investigated within genes (promoters, exons and introns), intergenic regions and transposable elements (TEs) (Figure 1c and d). More than half (52%) of the SMPs in CpG context fell into the intergenic regions, 7% in promoters, 28% in exons and 13% in introns (defining 48% within genes). In the CHG context, the number of SMPs in intergenic regions decreased to 37%, while we observed an enrichment in exons (35%) and introns (21%) and only 5% in promoter regions. Similarly, in the CHH context 5% of the SMPs were located in promoter regions, 48% in genes (27% in exons and 21% in introns) and the remaining 47% in intergenic regions (Figure 1d). TEs showed a higher level of methylation in comparison to genes in all the three contexts (Figure 1c and d). While promoter and genic regions displayed similar levels of methylation in non-CpG contexts (*i.e.* CHG and CHH), gene bodies were more methylated than promoter regions in the CpG context (Figure 1c and 1d). Overall, 93% of poplar genes (40,091 out of 42,950) and 83% of annotated TEs families (6,131 out of 7,386) were covered by bisulfite-converted sequence reads in the investigated populations (Figure S2).

**Figure 1:**
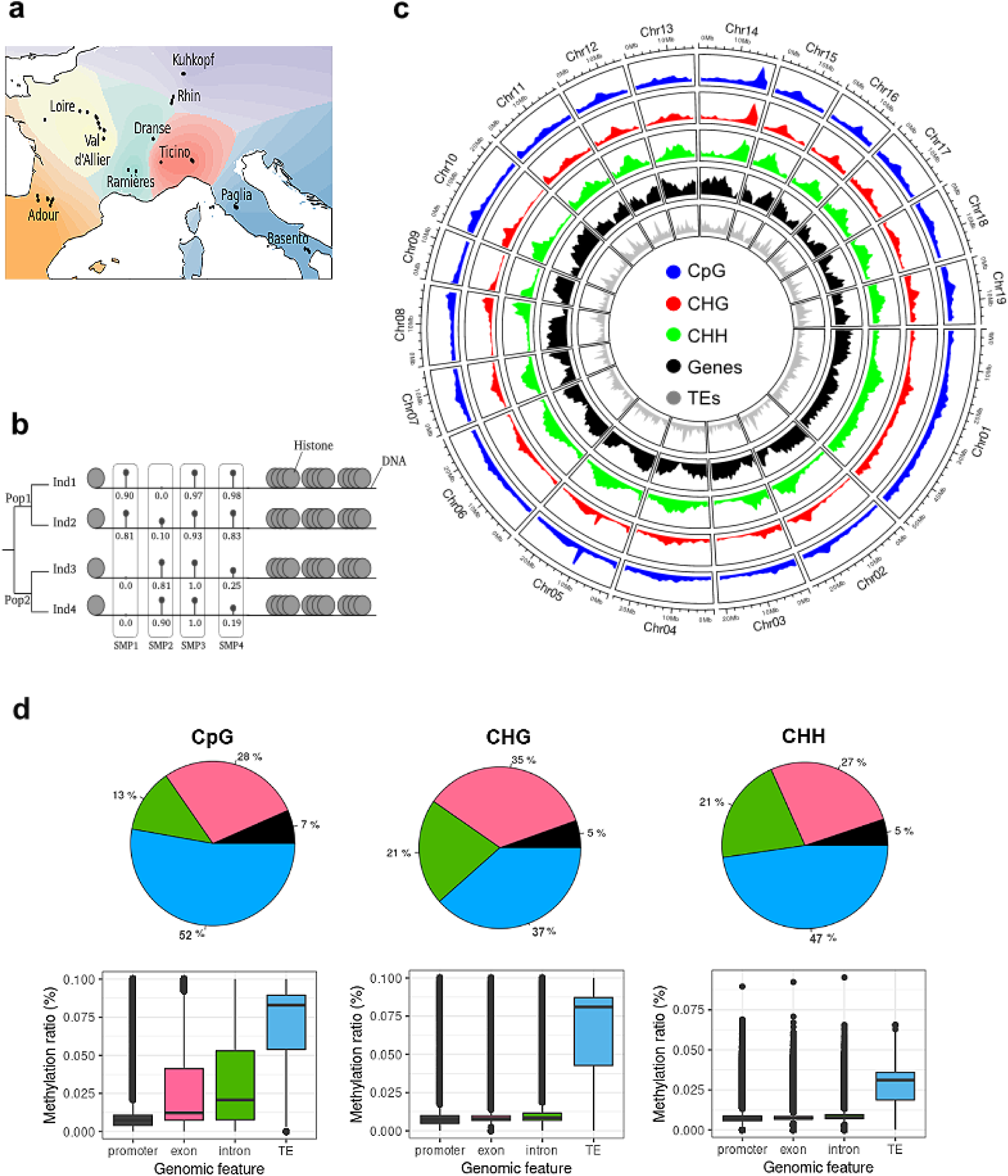
Methylation landscape in poplar. **A**, Geographical distribution of the 241 genotypes (black dot) from natural populations of *Populus nigra* representative of 6 genetic clusters (one by color) from a model-based ancestry estimation in ADMIXTURE program. **B**, Schematic representation of the SMPs (Single methylation polymorphisms). Numbers represent methylation ratio (mCs/total C) in a specific genomic position; Pop for Population; Ind for Individual. **C**, Circos plot of the distribution of DNA methylation in CpG, CHG and CHH contexts along genes and TEs for the 19 chromosomes of poplar. Methylation data are plotted for 1Mb windows with a minimum of methylation at 25% and 10X of coverage. CpG methylation density in blue; CHG methylation density in red; CHH methylation density in green; Gene density in black; TEs density in red. **D**, Annotation and methylation level (in percentage) of the SMPs (Single Methylated Polymorphism) in genomic features, i.e., in promoters (black), exons (red), introns (green) and intergenic or TEs (blue) for the three methylation contexts, CpG, CHG and CHH. Methylation analysis are shown for the Loire population.

### DNA methylation variation at the macro-evolutionary scale

It has been reported that whole genome duplications (polyploidizarions) drive (epi)genome reprogramming (Bellec et al. 2023). Within the angiosperms, poplar evolved from an Ancestral Eudicot Karyotype with 7 prochromosomes (AEK7) that went through whole genome triplication (*γ* WGT, ≈120 Mya ago) leading to AEK21 with 21 prochromosomes at the basis of any modern Eudicot species (Figure 2a), Murat et al. 2017. Within the Salicaceae botanical family, the modern poplar genome (19 chromosomes) derived from a n=12 ancestor (inherited from the AEK21 with 6 chromosomal fissions and 15 fusions) that has been duplicated (≈60 Mya WGD, to reach a n = 24 intermediate) followed by four chromosomal fissions and nine fusions (delivering a chromosomal equation of 19 = [21 + 6 - 15] x 2 + 4 - 9; Murat et al. 2015). Comparison of gene order between the inferred Eudicot ancestors (AEK7 and AEK21) and the modern poplar genome allowed to investigate the impact of ancient (ancestral γ WGT, ≈120 million years ago) and more recent (poplar specific WGD, ≈60 million years ago) polyploidization events on genome regulation (Figure 2). Polyploids are derived from two parental sub-genomes that were merged in the same nucleus and evolved through inversions, translocations, chromosomal fusion, and fission processes, leading to possible sub-genome differentiation, referred to as subgenome dominance (Bellec et al. 2023). The subgenome dominance is manifested by the differential retention of the ancestral gene content leading to least-fractionated regions (LF) and most-fractionated regions (MF) compartments in the modern poplar genome. Thus, we developed a synteny-based approach to detect genome compartments that underwent different fractionation after the ancestral (γ) WGT and poplar-specific WGD events (Figure 2). Overall, 6,442 genes were identified in LF compartments, and 3,356 genes in MF compartments (Figure 2a, b). GO enrichment for molecular functions showed that LF related genes were enriched for binding process (protein binding, mRNA binding, etc.), signaling (metal ion) and transferase activity, whereas genes in MF fractions were mostly enriched in binding and transcription factor activity (Figure S3). While duplicated genes may return into single copy (defining LF and MF compartments), some of the duplicates remain conserved as pairs in the modern genome following polyploidization events. A total of 2,020 duplicated genes inherited from the poplar-specific WGD were identified and found to be functionally enriched in binding, transcription factor activity and signaling processes (Figure S3).

**Figure 2:**
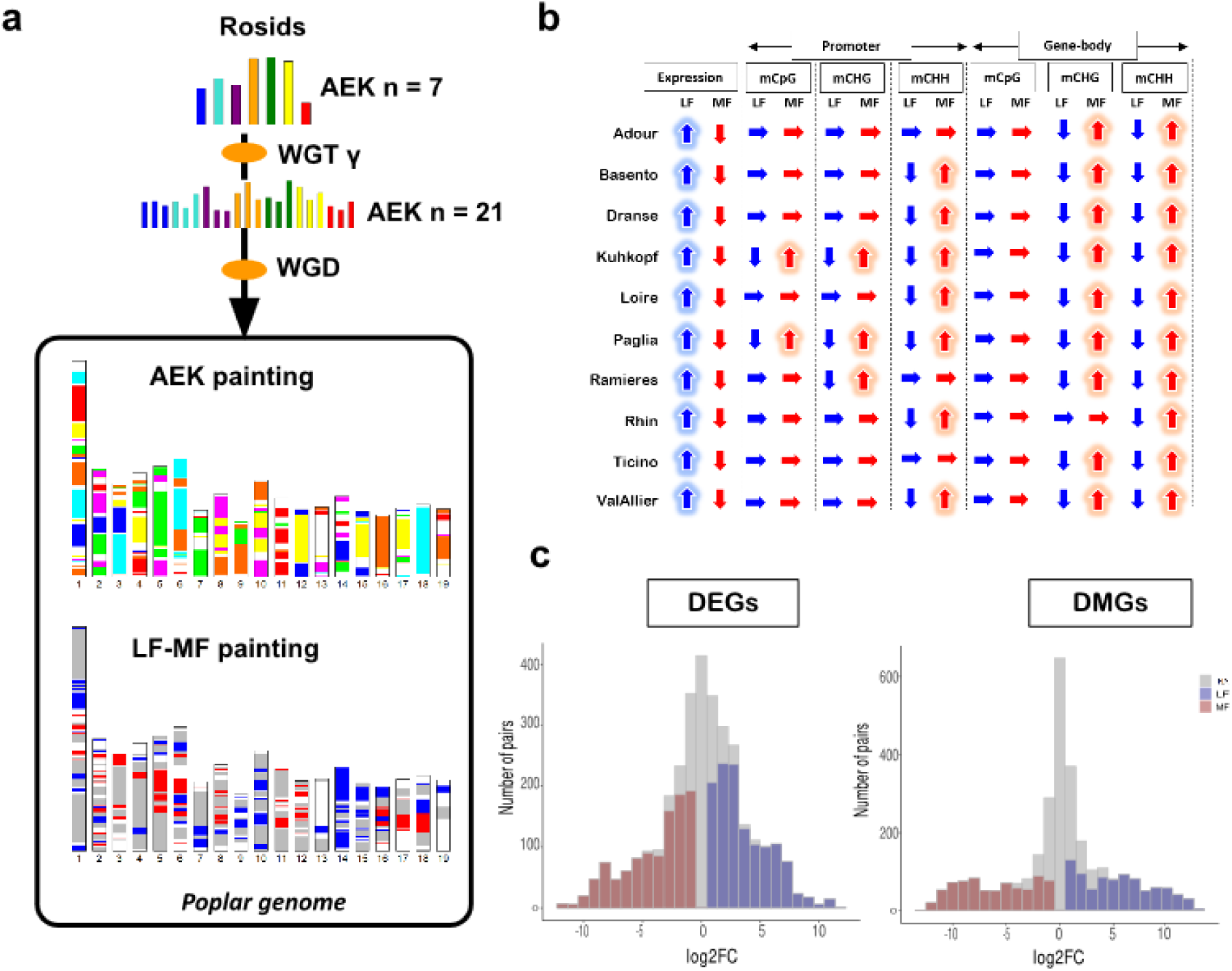
Genome evolution following ancestral polyploïdizations. **A**, Poplar genome from rosid ancestors and structural variations. Rosid ancestors with 7 reconstructed proto-chromosomes experienced a whole genome triplication (WGT) event to give rise to AEK21 with 21 proto-chromosomes. Salicales order then went through a second round of whole genome duplication (WGD) events followed by structural rearrangements ending with 19 chromosomes in poplar. Ancestral gene loss leads to different compartmentalization, LF (least-fractionnated, blue) and MF (most-fractionated, red). **B**, Evolutionary trajectory between LF (blue) and MF (red) genes in terms of methylation (promoter and gene body for CpG, CHG and CHH) and gene expression. Upper arrows indicated higher expression or methylation levels; Down arrows indicated lower expression or methylation levels; Horizontal arrows indicated no bias in expression or methylation levels. **C**, Differentially expressed genes (DEGs) and differentially methylated genes (DMGs) in all populations between duplicated gene pairs from the Salicales specific WGD. Blue for upregulated / hypermethylated LF-genes; Red for downregulated / hypomethylated genes in MF; Grey for no DEGs or DMGs between the duplicated genes in LF and MF. LogFC for log fold change. DEGs and DMGs are considered when FDR < 0.05 and logFC > 1.

To investigate the impact of polyploidization in shaping (epi)genome regulation over a macro-evolutionary scale, we compared expression and methylation profiles between genes in LF or MF fractions. No difference was observed for methylation in gene bodies (exons + introns) in CpG context between LF- and MF-located genes. However, in non-CpG contexts, MF-genes displayed higher methylation levels in comparison to genes located in LF compartments. The methylation level in gene promoters showed a distinct pattern. While no clear bias was observed for CpG and CHG contexts, MF-located genes appeared more methylated in their promoters than genes located in LF fractions (in seven out of the ten investigated populations) in CHH context (Figure 2b). Conversely, the genes located in the LF fraction appeared to be more expressed than genes in the MF fraction, displaying an antagonist pattern with DNA methylation (Figure 2b). Overall, regarding the ancestral (γ) triplication shared within the Eudicots, the LF compartment (where ancestral genes are preferentially retained) showed higher gene expression and lower methylation for promoters in CHH context and in gene bodies in CHG and CHH contexts, compared to genes located in MF compartments (where ancestral genes have been preferentially lost). To better understand the effect of polyploidization on gene regulation, we further investigated methylation and expression profiles between duplicated genes inherited from the recent poplar specific WGD. Interestingly, about ¾ of duplicated genes showed gene expression differences between the two copies of a pair. One of the copies was more expressed than the duplicated counterpart, and most of these over-expressed copies are located in the LF fractions of the genome. For the remaining ¼ of the duplicates, the two copies displayed the same expression level (Figure 2c). Similarly, about half of the duplicated genes showed methylation differences between LF and MF copies, while half of them expressed the same methylation level (Figure 2c). Furthermore, 72% of the differentially methylated gene pairs exhibited also expression differences (Figure S4). Overall, at the macro-evolution time scale, since the ancestral polyploidization events dating back to ≈60 and ≈120 million years ago, polar experience intense structural (gene loss) and regulation (expression and methylation) reprogramming. Genes from the MF subgenome were found to be more methylated (especially in non-CpG contexts) and less expressed compared to genes located in the LF fraction.

### DNA methylation variation at the micro-evolutionary scale

We then assessed the methylation landscape at a more recent evolutionary scale across ten populations, using a methylation threshold above the mean methylation in each context separately (*i.e.* a genomic feature was considered methylated when its methylation ratio was above the mean methylation observed from the ten populations, Figure 3a). We then reported between 20,433 and 21,321 genes/TEs methylated in the CpG context in the ten populations, between 9,324 and 9,785 in CHG and between 10,697 and 14,667 in CHH. Exploiting the concept of pan-genome (representing the entire set of genes within a species), consisting of a core genome (containing genes shared between all individuals of the species) and the ’dispensable’ genome (containing genes specific to individuals of the species), the pan-methylome consisted of an entire set of 25,485, 13,715 and 29,651 methylated genes and TEs in CpG, CHG and CHH contexts respectively, across the ten populations. The core methylome (*i.e.,* genes/TEs methylated in the ten populations) also varied between the three methylation contexts. The core methylome represented 66.6%, 52.5% and 25.4% of the pan-methylome in the CpG, CHG and CHH context, respectively, suggesting more stable CpG and CHG methylation between populations compared to CHH methylation (Figure 3a).

**Figure 3:**
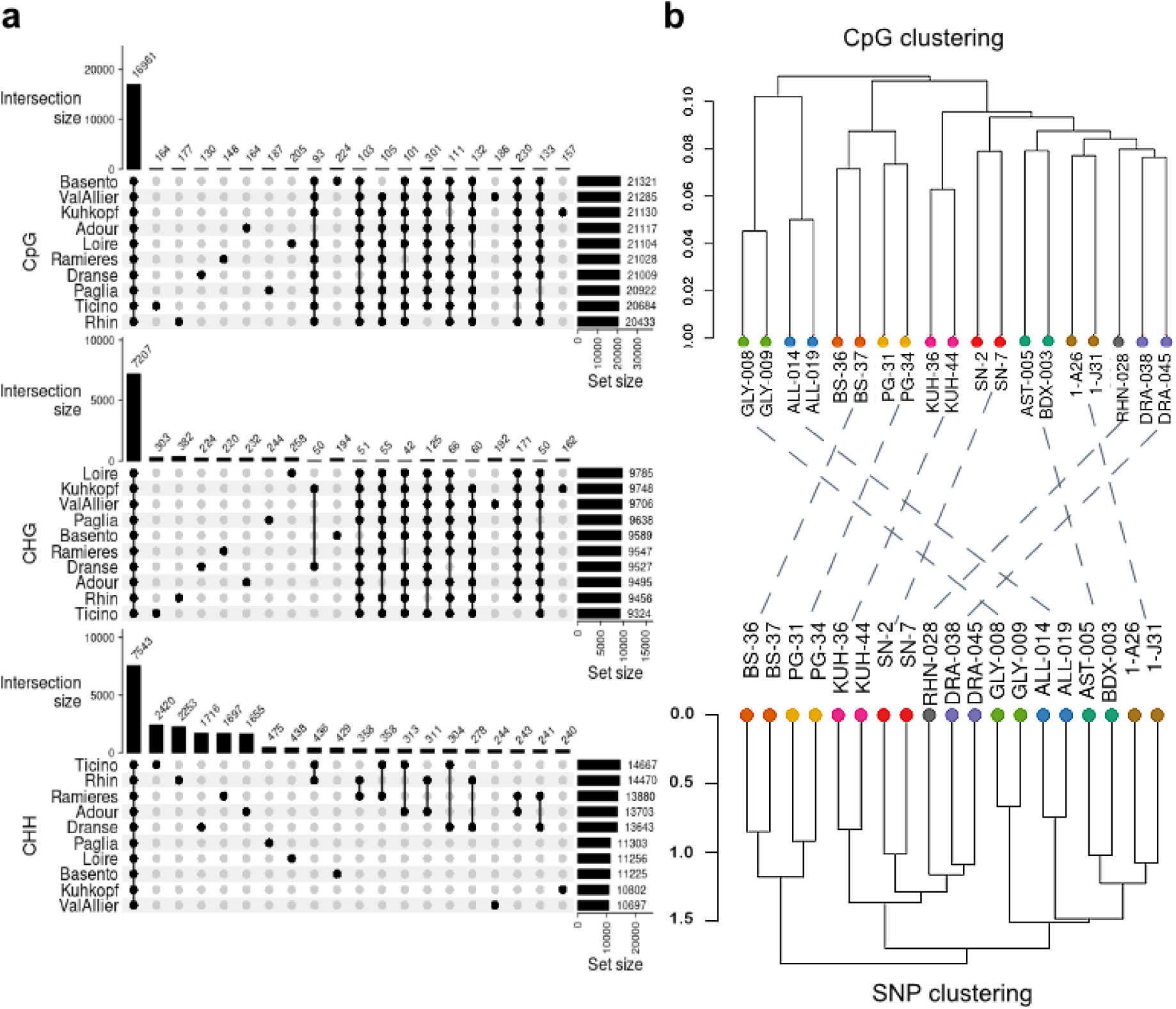
Phylo-epi/genomics in *Populus nigra*. **A**, Pan methylome of the ten natural populations in CpG, CHG and CHH contexts. The intersection size represents the number of common features between populations and the set size (horizontal bars, in right), represents the number of methylated features (genes and TEs) in each population. **B**, Phylo-epigenomic and phylogenetic tree reconstruction for DNA methylation (CpG) and genetic (SNPs) markers. The epigenetic trees were realized with genome-wide SMPs filtered with SNPs data, coverage (>7X) and tolerating 30% of missing data. The genetic tree was done with all the genome-wide SNPs without any missing value and with a minor allele frequency above 5%.

Furthermore, we investigated population relatedness solely based on available SMPs in all the investigated populations, through a phylo-epigenomic approach. Strikingly, the methylation clustering (strictly excluding SNPs) in CpG and CHG contexts recovered the geographical partitioning of the 10 natural populations, while such structure signal was partially lost in the CHH context (Figure 3b, Figure S5). Namely, CpG and CHG methylation clusterings fit the geographical origin of the populations defining three sub-groups. In addition to the phylo-epigenomic clustering we classically assessed the genetic structure (from SNP data) of the ten *P. nigra* populations. The hierarchical ascendant clustering on the genomic relationship matrix revealed three different sub-groups. The first sub-group was composed of two Italian populations (Basento and Paglia). The second sub-group consisted of four populations, two originating from France (Dranse and Rhine), one from Italy (Ticino) and one from Germany (Kuhkopf). The third sub-group was made of four French populations (Ramieres, Adour, ValAllier and Loire), Figure 3b. The genetic structure of the considered populations was consistent with their geographical origins and relatedness across west-east (sub-groups 1 and 3) and north-south (subgroups 1 and 2) axes (Figure 1a). Interestingly, the genetic structure was similar to the methylation clustering (CpG and CHG) suggesting that both genetics (DNA polymorphism) and epigenetics (DNA methylation) may act as markers of population differentiation.

### Role of DNA methylation in local adaptation

To unravel the biological functions driven by genes most differentiated at the epigenetic levels between the geographical origins, we focused on SMPs located in genes (including promoters) that best support the phylo-epigenomics and genetic structure of the ten studied populations (Figure S6a). Overall, 69,189, 18,523 and 11,282 SMPs in CpG, CHG and CHH contexts respectively, show significant association with the genetic structure of the populations (Figure 4a, Figure S6b). Based on a pcadapt analysis, we identified 2,241 (in CpG), 940 (in CHG) and 389 (in CHH) genes whose methylation profiles contribute the most to the differentiation of the natural populations (Figure 4b, Figure S6c). To obtain a more robust set of candidate genes supporting geographical differentiation, we considered an intersection of the previous pcadapt-derived genes, and those differentially methylated across the populations, resulting in 271 non-redundant genes (Figure 4c). Interestingly, many of these genes were enriched in functions related to disease resistance (34 TIR-NBS-LRR class, *i.e*. R genes), immune response (8 genes), hormonal and stress response (3 genes, MeSA, MeJA and DRY2). Methylation level of the disease resistance genes showed wide diversity among *P. nigra* populations (Figure 4d, Figure S7). For instance, R-genes in Basento, Paglia and Ticino populations from the southern range (Italy) were weakly- or un-methylated, which contrasted with the northern populations such as Loire and ValAllier (France).

**Figure 4:**
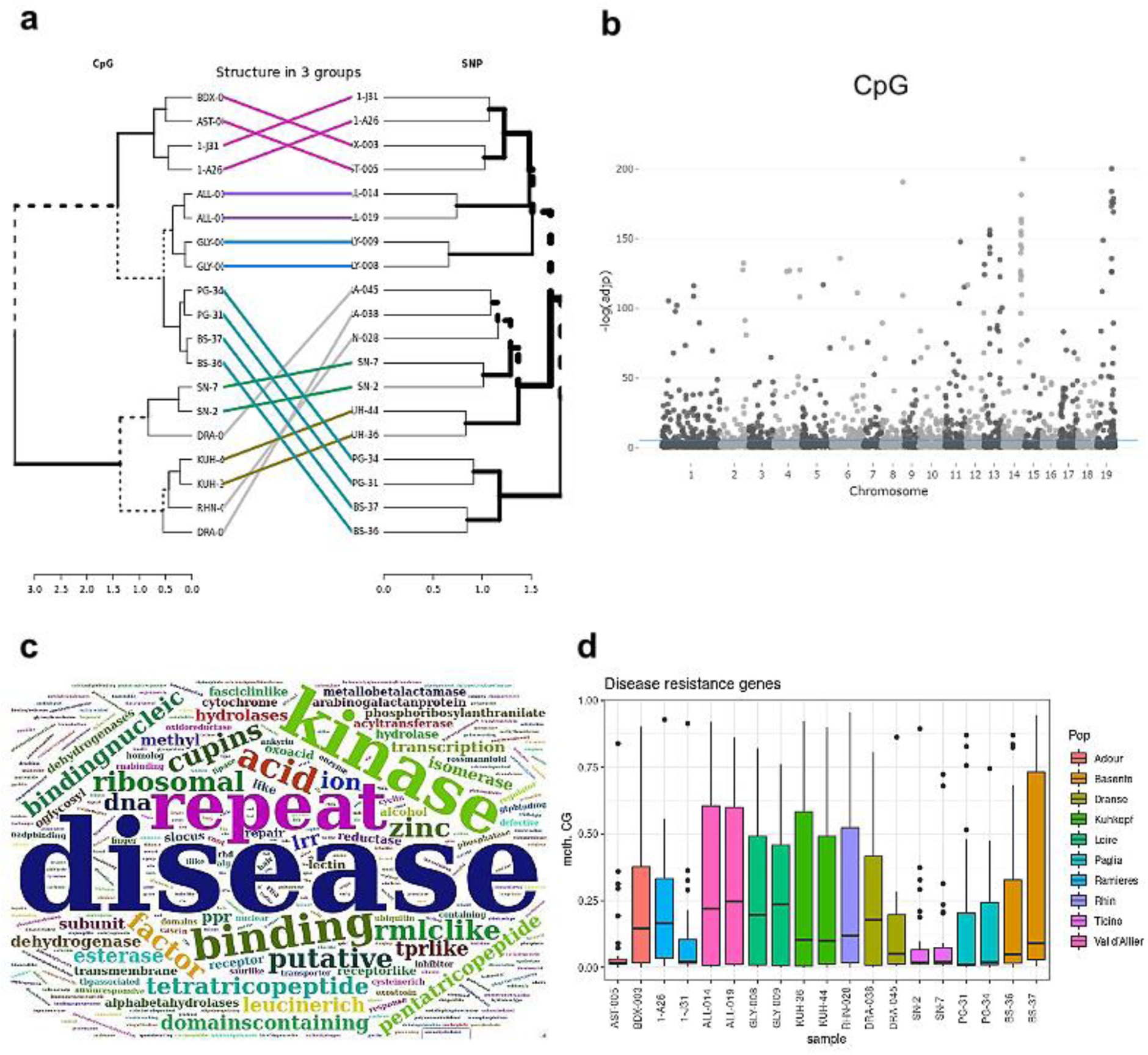
Identification of genomic markers of local adaptation and/or drift. **A**, Comparison between the phylo-epigenomic tree in CpG built on differentially SMPs between the three sub-groups and the phylogenetic tree. **B**, Manhattan plots of obtained epigenetic pcadapt gene markers in CpG context with adjusted p-values (FDR, blue line). Markers above the blue line (FDR = 0.05) are considered as putative markers of local adaptation. **C**, WordCloud of identified putative genes involved in local adaptation. **D**, CpG Methylation diversity of disease resistance (R) genes in the 20 black poplar trees. The color code represents the population groups.

### Interplay between DNA methylation and gene expression

To unveil the possible link between DNA methylation and gene expression at the population level, we compared methylation in % (standard normalization) or in rbd (Bellec et al. 2023) to gene expression (in TPM) (Figure S8a). When assessing the link between DNA methylation (in %, CpG) and gene expression at the whole genome level, no clear pattern could be established (Figure S8b, spearman correlation *r* = -0.095, *p-value* < 2.2e-16). However, were able to clearly distinguish three groups of genes combining distinct patterns of expression and methylation. Highly expressed genes in populations displaying low or no methylation (hereafter called Hypo/Up, for hypomethylated and up-regulated genes), highly methylated and lowly expressed genes (hereafter called Hyper/Down, for hypermethylated and down-expressed) and genes that are both methylated and expressed (Figure S8b). Given this result obtained with methylation expressed rbd (methylated reads are weighted with the ratio between methylated and un-methylated reads allowing to focus only on reads supporting the methylation status), we developed a different approach to study the relationship between gene expression and DNA methylation (in %, the standard methylation normalization) by creating methylation ratio quantiles with 10% increment (Figure 5a and b). Comparison between gene expression and promoter methylation percentage (in quantile) clearly showed that the expression of most genes is negatively correlated to increase in methylation level in each methylation context (Figure 5a, Figure S8c). Using the same approach, we investigated the relationship between methylation in gene bodies and gene expression. The expression of genes increased gradually with the methylation level in the CpG context before a strong decrease when methylation reached ∼60% (Figure 5b). However, gene body methylation in CHG and CHH contexts displayed a similar pattern observed for promoter methylation (Figure S8d). Overall, it appears that DNA methylation at the population level affects gene expression in a discontinuous manner, having no discernible effect at low-level changes but probably leading to silencing when a certain threshold is reached, either in the promoter or the gene body, pointing out a dosage effect of methylation on gene expression.

**Figure 5:**
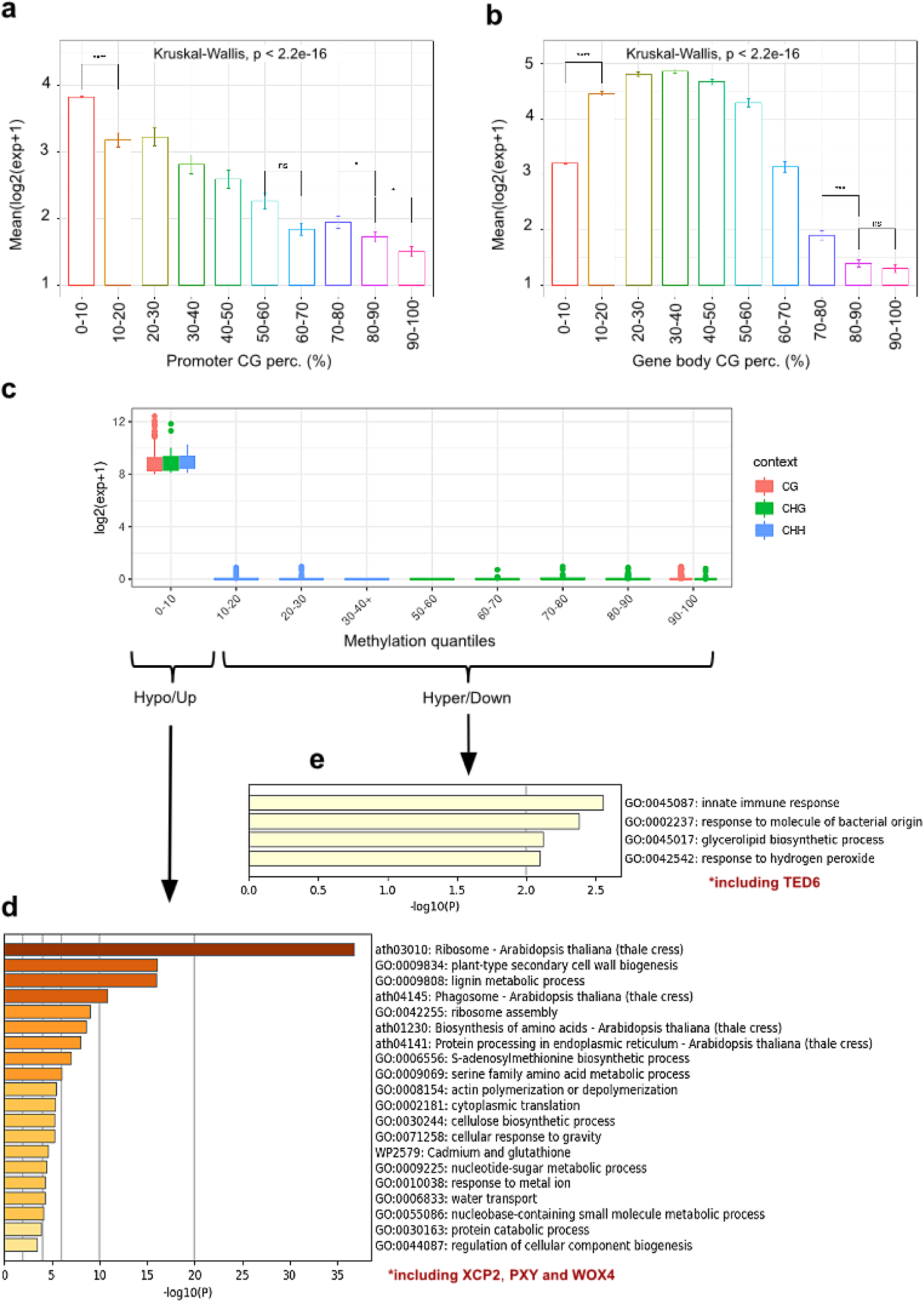
Regulation of gene expression by DNA methylation. **A**, Relationship between promoter DNA methylation in CpG context and gene expression. Methylation data are splitted into 10 quantiles with each quantile capturing 10% of the methylation level. Expression data are shown as the mean expression of genes in log2. **B**, Relationship between gene body DNA methylation in CpG context and gene expression. Methylation data are splitted into 10 quantiles with each quantile capturing 10% of the methylation level. Expression data are shown as the mean expression of genes in log2. **C**, Identification of outlier genes between promoter DNA methylation and gene expression (i.e. Hypo/Up and Hyper/Down). Methylation (x axis) is represented by quantiles with each quantile capturing 10% of the methylation level and gene expression (y axis) by logarithm. Red for CpG context, green for CHG context and blue for CHH context. Hypo/Up for hypomethylated and overexpressed genes and Hyper/Down for hypermethylated and downregulated genes. **D**, GO annotation using biological process terms for Hypo/Up genes. **E**, Functional annotations for Hyper/Down genes using biological process terms.

Which physiological traits can be driven through expression-methylation interplay? To address this question, we then focused on genes showing the most contrasted methylation and gene expression profile among populations, *i.e.* Hyper/Down and Hypo/Up genes (Figure 5c). Such genes highlight that DNA methylation may control the tissue-specific expression (from cambium and xylem in the current experiment) of targeted genes. Functional annotation of Hypo/Up genes (*i.e.* weakly- or un-methylated while highly expressed) showed enrichment in functions related to ribosome process (75 genes of housekeeping functions), cell wall and lignin metabolic process (53 genes involved in cambium differentiation), etc. that are essential functions for cell growth and differentiation (Figure 5d). Namely, *XCP2* (XYLEM CYSTEINE PEPTIDASE 2), *WOX4* (a WUSCHEL-related homeobox gene family member) and *PXY* (PHLOEM INTERCALATED WITH XYLEM) involved in xylem and phloem differentiation, were found to be constantly expressed and not methylated. Similarly, GO annotation of Hyper/Down (*i.e.* highly methylated while lowly- or not-expressed) genes revealed enrichment in immune response and glycerolipid biosynthetic process (Figure 5e) with two *MYB* transcription factors but also an important key gene for cambium formation. Namely, *TED6* (tracheary element differentiation-related) involved in differentiation of xylem vessel elements by promoting secondary cell wall formation is apparently constantly silenced by high DNA methylation level. Our data suggest that fine-tuned interplay between omics (expression/methylation) data may control the physiological development and differentiation of the considered tissues (cambium and xylem).

## DISCUSSION

Contrary to model species (*A. thaliana*) or crops (tomato and rice) where most of epigenetic studies have been conducted in herbaceous and annual plants so far, poplars are woody long-lifespan trees that are repeatedly exposed to environmental challenges over decades. Therefore, trees have developed various mechanisms enabling them to adapt and to survive. Several previous studies in annual plants found correlations between methylation patterns and habitat or climate in different plant species (Kawakatsu et al. 2016; Xu et al. 2020; Galanti et al. 2022), but there is lack of evidence of such phenomenon, if existing, for perennial species. In this study, we present the analysis of natural populations of black poplars (*Populus nigra*) sampled from four distinct geographic origins (France, Italy, Germany and Netherlands) and grown in a common garden in France. We focus on cambial tissues, an important functional trait for forest trees responsible of wood formation, to assess the evolutionary and functional impact of epigenetic variations at macro- (past polyploidization events) and micro- (geographical origins) scales.

### DNA methylation as a stable epigenetic mark in tree populations

How DNA methylation is acting on genomes of different tree populations? DNA methylation has been studied in many plant species, showing wide diversity in terms of methylation level and pattern (Bartels et al. 2018; Niederhuth et al. 2016). Methylation diversity is observed for all the three methylation contexts (CpG, CHG and CHH) even though CpG methylation (the predominant type) is more stable across species (Niederhuth et al. 2016). Here, we assessed DNA methylation at the population level in black poplars. Methylation level was quite stable for CpG and CHG contexts across populations, and more variable in the CHH context. About half of methylated cytosines were located in genomic features, including promoter regions. Transposable elements appeared to be more methylated in all three methylation contexts compared to genic features, a general trend observed in many species (Cao et al. 2021; Zhang et al. 2018; Zhang et al. 2006). The methylation of TEs was much higher in CpG and CHG contexts, a feature detected in almost all studied plant epigenomes (Bräutigam and Cronk 2018; Lang et al. 2018; Zemach et al. 2010). The distribution of DNA methylation across the investigated genomic features were also significantly different between the methylation contexts. While in the CpG context, genic regions are more methylated than promoters, they display similar methylation levels in the non-CpG contexts. The increased CpG methylation level in genic regions could be related to DNA methylation-driven silencing of intron-located repetitive elements (Cao et al. 2021). Moreover, the pan-methylome consisted of 25,485, 13,715 and 29,651 methylated genes and TEs in CpG, CHG and CHH contexts respectively across the studied *P. nigra* populations when applying a methylation threshold above the mean value observed in the populations in each context. Specifically, 66,6%, 52.4% and 25.4% of poplar genes/TEs were constantly methylated (core methylome) in CpG, CHG and CHH contexts, respectively. Although DNA methylomes are available for many plant species, integrative analysis of the DNA methylome profiles in a pan-methylome manner (between individuals-tissues of the same species) has been little studied in plants. In humans, the comparison of DNA methylation patterns in different cancer types has provided novel insights into alterations that contribute to cancer development (Shimizu et al. 2021; Liu et al. 2018). Here, we show that 16,961, 7,207 and 7,543 genes and TEs are methylated in all 10 black poplar trees, in CpG, CHG and CHH context, respectively. Only the CHH methylation context expresses wide diversity between the different populations and highlights the conservative nature of DNA methylation in *P. nigra* populations (core methylome greater than 50% of the pan methylome in CpG and CHG). Overall, these results provide a comprehensive analysis of genome-wide DNA methylation across natural populations of a keystone species of the riparian forest ecosystems, highlighting the conservative nature of methylation patterns across the populations especially in CpG and CHG contexts.

### DNA methylation and gene expression reprogramming following ancestral polyploidization events

Do epigenetic modifications provide a signal of long-term evolution following polyploidization events? Polyploidization (or WGD) has occurred frequently during plant genome evolution and represents an evolutionary driving force which provides extra genetic material to be specialized for phenotypic diversification, adaptation, and survival. Such an event is followed by a diploïdization (or extensive fractionation) process reverting the polyploids to diploid status through gene number reduction (Van de Peer et al. 2021; Soltis et al. 2015; Wendel, 2015; Murat et al. 2012). Ancestral gene losses during the diploidization process (accompanied by genomic structural rearrangements) may lead in some cases to subgenome dominance in the form of biased compartmentalization, with dominant (retaining more ancestral genes) and sensitive (retaining fewer ancestral genes) subgenomes, so called least-fractionated (LF) and most-fractionated (MF) regions (Alger and Edger, 2020; Cheng et al. 2016). We conducted a comparative analysis between Eudicot ancestors (AEK7 and AEK21) and the modern poplar genome to detect fractionation bias in ancestral gene retention after polyploidization events. Instead of focusing on the specific *Salicaceae* WGD which was inferred to have occurred around 60 million years ago (Dai et al. 2014) and where no subgenome dominance has been reported (Liu et al. 2017), we investigated the evolutionary impact of the γ triplication event shared by all the rosids (∼120 mya) in terms of gene loss, DNA methylation and gene expression. Hence, 15% of the annotated poplar genes could be attributed to the dominant-LF compartment and 8% to the sensitive-MF compartment. Functional annotation of genes in the dominant-LF fraction showed enrichment in binding and signaling process, whereas MF-genes are more involved in transcription factor activity as previously reported (Blanc and Wolfe, 2004; Freeling, 2009; Hao et al. 2021). The LF compartment showed a higher gene expression and lower methylation levels for promoters in the CHH context and for genic regions in CHG and CHH contexts, compared to genes located in the MF compartment. This has also been reported in cotton, maize and *Brasicaceae* where the dominant-LF subgenome appeared to be more expressed and less methylated (Renny-Byfield et al. 2017; Woodhoouse et al. 2014; Schnable et al. 2011). Besides subgenome dominance, we further investigated the regulation plasticity of duplicated genes (that remain conserved as pairs after the diploidization process) derived from the specific *Salicaceae* WGD. It has been proposed that duplicated genes, initially having identical sequences and functions, tend to diverge in regulatory and coding regions, which may change their expression pattern or lead to the acquisition of new functions (Blanc and Wolfe, 2004; Xu et al. 2012). In poplar, about ¾ of the duplicated genes showed gene expression differences and are preferentially located in the dominant-LF fraction of the genome. The number of duplicated pairs with expression differences vary widely between species (Zhao et al. 2017; Schnable et al. 2011; Throude et al. 2009; Yim et al. 2009). Similarly, about half of duplicated genes in poplar showed methylation differences. Keller and Yi (2014) highlighted that for a majority of duplicate gene pairs in humans, a specific duplicate partner is consistently hypo- or hypermethylated across highly divergent tissues. Analysis of duplicated gene pairs in cassava showed that gene body methylation and gene expression have co-evolved within a short evolutionary timescale (Wang et al. 2015; Wang et al. 2017). Here, we found that 72% of differentially expressed genes showed methylation differences. Overall, after past polyploidization events, poplar showed divergent evolutionary patterns reflected by subgenome dominance, gene expression and epigenetic differences.

### DNA methylation as a marker of population differentiation alike genetic markers

To what extent do epigenetic marks differentiate populations from different geographic origins? Besides genetic variation, natural plant populations usually also harbor epigenetic variation. This idea arose by the fact that DNA methylation variation in natural plant populations is often non-random and geographically structured (Dubin et al. 2015; Garino et al. 2015; Gugger et al. 2016; Kawakatsu et al. 2016; Galanti et al. 2022). Using a phylo-epigenomic approach, we clearly established that DNA methylation, in the same manner as genetic markers, differentiate population structure according to their geographic origins. This result suggests that DNA methylation, at least for CpG and CHG contexts, could be used for genotype and/or population differentiation just like SNPs. Thus, epigenetic mechanisms could represent an additional layer of heritable phenotypic variation since these modifications can be inherited through generations and possibly explain a part of the missing heritability of Mendelian traits (Maher, 2008; Becker et al. 2011; Heckwolf et al. 2020; Noshay and Springer, 2021; Sammarco et al. 2022). Within natural populations of oak in Southern California, Platt et al. (2015) found patterns of genetic and epigenetic (mainly CpG methylation) differentiation indicating that local adaptation is operating on large portions of the oak genome. Methylation in the CpG context is more frequent but also more abundant at the gene level, suggesting that it may have more adaptive ‘power’ compared to non-CpG methylation. This hypothesis is supported by Gugger et al. (2016) who identified variations in CpG methylation linked to climatic variations at or near genes, suggesting a direct relationship between DNA methylation in CpG and local adaptation. However, Platt et al. (2015) did not establish a link between methylation in CHG context and local adaptation and suggested that CHG methyl-polymorphisms are not playing a significant role. In the current study we clearly established that CHG methylation clustering fits the geographic distribution of the natural populations and is similar to their genetic clustering. Nonetheless, CHG methylation levels vary more widely across species compared to CpG methylation (∼10.0% in *A. thaliana* against ∼26.8% in *P. trichocarpa*, Barlets et al. 2018) and may therefore have different impacts depending on the species. In our populations, the overall methylation level in CpG and CHG contexts was quite high, ranging between 32.1% to 35.4% for CpG and between 20.0% to 21.9% for CHG, depending on the considered population. These results may suggest that the geographic structure of the population origins is genetically and more interestingly epigenetically marked and that DNA methylation needs to be considered as a key marker of species micro-evolution (over millennia of evolution) in complementing the Mendelian principle of genetic inheritance as proposed by Jablonka (2017). However it is not clear whether such results also illustrate epigenetic drift or to what extent epigenetic variations follow a demographic structure determined by genetic markers (Lamka et al. 2022). Galanti et al. (2022) showed that in *Thlaspi arvense*, natural epigenetic variations are significantly associated with both genetic variation and environment of population’s origin, and that the relative importance of the two factors strongly depends on methylation contexts, with environmental variations being higher in non-CG contexts. Hence, the CHG population structure reported here may support the hypothesis of a role of DNA methylation in population differentiation. While some publications mention genetic control over epigenetic variations (Becker et al. 2011; Dubin et al. 2015; Sow et al. 2018b, Alvarez et al. 2021), epigenetics could also accelerate mutational dynamics (Ossowski et al. 2010; van der Graaf et al. 2015 Johannes, 2019; Zhou et al. 2020) and therefore could mimick the geographical structuring of both phylogenetic and phylo-methylomics structures.

### DNA methylation reshapes gene expression dynamics among populations

While DNA methylation is quite often associated with regulation of gene expression, it is more likely that instead of being a simple “on-off switch”, DNA methylation has a nuanced impact on the expression of genes according to targeted genomic features and contexts (Zhang et al. 2006; Niederhuth and Schmitz, 2017; Bewick and Schmitz, 2017). However, even when DNA methylation was evaluated per genomic features (*i.e.* promoter and genic regions), we observed poor correlation between DNA methylation and gene expression on the whole genome level among populations. This result is in contrast with common knowledge that DNA methylation in promoters is associated with gene silencing (Kon and Yoshikawa, 2014; Nuo et al. 2016; Bewick and Schmitz, 2017; Ma et al. 2020), but consistent with the findings of Li et al. (2012), who showed that methylation in promoter regions repressed only a few heavily methylated genes in rice. In order to scrutinize in more depth, the relationship between DNA methylation and expression, we categorized genes according to quantiles of their methylation level. We first assessed the link between promoter methylation and gene expression and revealed a significant negative correlation with gene expression when promoters are highly methylated. The same pattern was reported during transgene inactivation in transgenic plants (Weinhold et al. 2013; Nuo et al. 2016). For methylation in genic regions, that has been positively correlated with gene expression (Ball et al. 2009; Bewick and Schmitz, 2017), we confirm, at the population level, the expected relationship in the CpG context, but only at low-to-moderate methylation levels. The positive correlation of genic methylation and expression is disrupted on the genome-wide scale by the fact that highly (methylation ratio above 60%) methylated gene bodies showed a strong negative correlation with gene expression. Overall, the link between DNA methylation and gene expression is more nuanced and complex than initially thought, with specificity on a gene-by-gene basis. Genome-wide patterns can only be disentangled when genomic features (i.e., promoter or genic regions) and sequence contexts are evaluated separately.

### DNA methylation regulation of key functional trait among populations

As long-lived species, trees are characterized by the expansion of disease-resistance (R) genes as a key process to face a wide range of biotic threats over their lifespans (Plomion et al. 2018). R genes are shown to enable plants to detect specific pathogen-associated molecules and initiate signal transduction to activate defenses (Hammond-Kosack and Jones, 1997). NBS-LRR (nucleotide binding site-leucine rich repeat) genes, a class of immune receptor, are well known to play fundamental roles in disease resistance (Dangl and Jones, 2001; Kong et al. 2020). The expression levels of plant NBS-LRR genes may be regulated by different mechanisms, including DNA methylation (Kong et al. 2020). In our study, we found that genes with different methylation profiles between natural populations of distinct geographical origins were enriched in functions related to disease resistance (TIR-NBS-LRR genes). While R-genes were weakly- or un-methylated in populations originating from Italy (Basento, Paglia and Ticino), they appeared more methylated in French populations such as Loire and ValAllier. This suggests that DNA methylation could represent a key mechanism to control the expression of R-genes in trees. Indeed, previous report in common bean has shown that most of R-genes are methylated, reminiscent of the DNA methylation pattern of surrounding repeated sequences (Richards et al. 2018a). As R-genes-triggered immunity can be associated with a reduction in growth and yield, so-called ‘fitness costs’, plants use an elaborate interplay of different mechanisms to control R-gene transcript levels to avoid the associated cost of resistance in the absence of a pathogen (Richards et al. 2018b). Hence, it appears that DNA de/methylation is required for the proper expression/silencing of defense related genes (Zeng et al. 2021). We also found that genes with different methylation profiles between populations are enriched in functions related to hormonal signaling. Such a relationship between DNA methylation and phytohormone related genes has been already reported in poplar (Lafon-Placette et al. 2018; Maury et al. 2019; Zhang et al. 2020; Sow et al. 2021). Similarly, *DRY2* (DROUGHT HYPERSENSITIVE 2, response to water deprivation, response to ethylene), another important gene involved in shoot growth, was found to be differentially methylated between the studied populations, suggesting variation in response to drought stress among the poplar gene pools (Wang et al. 2020).

We then investigated genes showing strong negative correlation between DNA methylation and gene expression, namely the Hypo/Up (weakly- or un-methylated and highly expressed) and the Hyper/Down (highly methylated while lowly- or not-expressed) genes, within the considered tissues, cambium and xylem. For Hypo/Up genes, we observed enrichment in housekeeping (ribosome activity) and cambium activity (e.g. in cell wall biosynthesis) related genes. Since housekeeping genes are required for the maintenance of basal functions for cell survival (Joshi et al. 2022), it is assumed that they are constitutively expressed and therefore depleted or lowly methylated to maintain their proper expression. Similarly, several genes essential for cambium differentiation were found weakly- or un-methylated and highly expressed. This is the case for (i) *XCP2*, a XYLEM CYSTEINE PEPTIDASE involved in programmed cell death of rays tylosis essential for heartwood formation (Avci et al. 2008; Nakaba et al. 2015; Zheng et al. 2015), (ii) *PXY*, a receptor-like kinase essential for maintaining polarity during plant vascular-tissue development (Fisher and Turner, 2007) and (iii) *WOX4*, a WUSCHEL-related homeobox gene family playing a crucial role in the regulation of cambium cell proliferation (Fisher et al. 2019). The vascular stem-cell tissue known as cambium generates phloem cells on one side and xylem cells on the other. While xylem is required for water transport, phloem is required primarily for the transport of photoassimilates (Sieburth, 2007; Etchells et al. 2015). Xylem and phloem form the vascular tissues of trees, an important functional trait for forest trees. It has been shown that in *pxy* mutants, the spatial organization of vascular development is lost and the xylem and phloem are partially interspersed (Fisher and Turner, 2007). Similarly, ectopic overexpression of *PXY* gene in hybrid poplar resulted in vascular tissue abnormalities and poor plant growth (Etchells et al. 2015). Moreover, *wox4* mutants exhibited reduced cell division activity in the cambial tissue (Fisher et al. 2019). Interestingly the two candidate genes, *PXY* and *WOX4* are known to act in the same signaling pathway, *WOX4* being downstream of the *PXY* receptor kinase to regulate stem cell proliferation (Etchells et al. 2013; Fisher et al. 2019; Hu et al. 2022). Recently, Dai et al (2023) using more than 20 sets of poplar transgenic lines have shown that *WOX4* system may coordinate genetic and epigenetic (histone marks) regulation to maintain normal vascular cambium development for wood formation.

For Hyper/Down genes, we observed an enrichment in innate immune response, suggesting a control of immune response genes by DNA methylation. In addition, *TED6* (Tracheary Element Differentiation-Related6) involved in xylem vessel differentiation (secondary cell wall) was found highly methylated and silenced. Transient RNAi of *Arabidopsis TED6* and 7 resulted in aberrant secondary cell wall formation of *Arabidopsis* root vessel elements (Endo et al. 2009). Moreover, it has been shown that a *rdd* mutant in *Arabidopsis* (defective DNA demethylation) with affected expression of many genes including *TED6* is impaired in tracheary element differentiation (Lin et al. 2020). Homology searches have identified *TED6/7*-like proteins only in the angiosperm lineage suggesting that the development of *TED6/7* proteins could have coincided with the emergence of the angiosperm lineage, and that they may have made key contributions to the evolution of water-conducting cells from tracheids to vessels (Rejab et al. 2015).

Overall, the current study shows that DNA methylation leaves genomic footprints recognizable at both macro- and micro-evolutionary scales and has a nuanced relationship with gene expression in poplar. In this study, genes with key functions for trees (disease resistance and wood formation) have been identified based on their DNA methylation patterns (on genome-wide or inter-population scales), suggesting a role of DNA methylation in their regulation. Our data showed that DNA methylation in poplar populations display natural variations, and may regulate fitness traits (disease resistance and wood formation). These results also highlight the need to take epigenetic markers into account in breeding strategies (Kakoulidou et al. 2021) together with genetic markers for both wood production and quality in the context of climate change that requires adaptation to biotic and abiotic constrains.

## EXPERIMENTAL PROCEDURES

### Sample collection and population structure

The initial experimental design consisted of 1,160 black poplar genotypes sampled from 14 river catchments of 4 European countries, Germany, France, Italy and Netherlands (Guet et al. 2015; Gebreselassie et al. 2017). The genetic diversity within this black poplar collection was previously characterized using 5,600 SNPs from a 12k Infinium array (Faivre-Rampant et al. 2016) and led to the definition of a subset of 241 genotypes representative of the genetic diversity while avoiding the widespread introgression from the cultivar *P. nigra* cv. Italica. These 241 genotypes originated from 10 river catchments (Chateigner et al. 2020). Population structure analysis was carried out on this restricted set of 241 genotypes with the same set of 5,600 SNPs and the model-based ancestry estimation in the ADMIXTURE program (Alexander et al. 2009) highlighted six genetic clusters which minimized the cross-validation error (Figure 1a).

### Genomic DNA extraction

For the present study, twenty genotypes (two genotypes per river catchment) were selected to be representative of the diversity of the black poplar collection. We sampled cambium and xylem on two biological replicates (clones) of each genotype, located in two blocks of a large common garden experiment at INRAE Orléans, France. The resulting 80 samples were further used to extract genomic DNA. DNA was extracPG_34_pe.bamted using a cetyl trimethy-lammonium bromide (CTAB) buffer according to Doyle & Doyle (1987). Extracted gDNA was then quantified using a Nanodrop spectrometer (Thermo Fisher Scientific, Waltham, MA, USA). An equimolar pool of the gDNA samples from the two individual clones and the two tissues (xylem and cambium, for wood formation) of each genotype was then performed. The gDNA pool for each of the 20 genotypes was sent to the CEA laboratory in Evry for both WGS (whole genome sequencing) and WGBS (whole genome bisulfite sequencing).

### Whole Genome Sequencing (WGS), read alignment and variant calling

Whole genome sequencing was performed by the ‘Centre National de Recherche en Génomique Humaine (CNRGH), Institut de Biologie François Jacob, CEA, Evry, France’. After a complete quality control, genomic DNA (1 µg) has been used to prepare a library for whole genome sequencing, using the Illumina TruSeq DNA PCR-Free Library Preparation Kit, according to the manufacturer’s instructions. After normalization and quality control, libraries have been sequenced on a HiSeqX5 platform (Illumina Inc., CA, USA), as paired-end 150 bp reads. One lane of HiSeqX5 flow cell was used for each sample, to reach an average sequencing depth of 30x for each sample. Sequence quality parameters were assessed throughout the sequencing run and standard bioinformatics analysis of sequencing data based on the Illumina pipeline was used to generate FASTQ files for each sample. We followed the bioinformatics pipeline described in Rogier et al. 2018 with small modifications. After a read quality control with *FastQC* v0.11.7 (Andrews, 2010), sequences were trimmed using the *Trimmomatic* tool v0.38 (Bolger et al. 2014). The adapter sequences were removed, the 9th first bases and the low-quality bases were trimmed based on a PHRED score below 20 and finally the reads with a length of less than 35 nucleotides were discarded. The mapping of the reads was performed using *BWA mem* v0.7.17 (Li, 2013) on the *Populus trichocarpa* v3.1 reference genome (Tuskan et al. 2006). Then, the *Picard Toolkit* v2.18.2 allowed the removal of the duplicated reads. Three caller programs were used to identify the variants: (i) *GATK* v4.0.11.1 (McKenna et al. 2010) using the *HaplotypeCaller* tool in single-sample calling mode followed by joint genotyping of the 20 samples; (ii) *FreeBayes* v1.2.0-2 (Garrison et al. 2012) in multi-sample calling mode; and (iii) *SAMtools* v1.8 (Danecek et al. 2021) using the *mpileup* tool in multi-sample calling mode followed by the *BCFtools* v1.8 (Li, 2011) call command. Finally, we considered only biallelic intra-nigra SNPs with quality threshold ≥ 30. Raw data were filtered with *VCFtools* v0.1.15 (Danecek et al. 2011) and only SNPs identified by at least 2 callers were selected to obtain the final set of SNPs.

### Whole Genome Bisulfite Sequencing (WGBS), read alignment and methylation calling and annotation

Whole genome bisulfite sequencing was performed using the Ovation Ultralow Methyl-seq kit (Tecan Genomics/Nugen, San Carlos, CA, USA, http://www.nugen.com/products/ovation-ultralow-methyl-seq-library-systems) following the published procedure (Daviaud et al. 2018). The workflow of the library preparation protocol follows a standard library preparation protocol, in which methylated adaptors are ligated to the fragmented DNA prior to bisulfite conversion. 200 ng of genomic DNA was fragmented to a size of approximately 200 base pairs (bp). Purified and methylated adaptors compatible with sequencing on an Illumina HiSeq instrument were ligated. The resulting DNA library was purified, and bisulfite converted. A qPCR assay determined the optimal number of PCR amplification cycles (between 10 to 15 cycles) required to obtain a high diversity library with minimal duplicate reads prior to the actual library amplification. The 20 samples to be sequenced were combined at equimolar quantities in two pools and the sequencing was performed in paired end mode (2 x 150 pb) on ten lanes of two Illumina HiSeq4000 flow cells to reach a minimal theoretical coverage of 30X for each sample. Fastq files from each sample were concatenated following sequencing.

The bioinformatic pipeline for DNA methylation analysis was executed on a Galaxy instance of the IHPE (Interactions Hôtes Pathogènes Environnements) platform (http://galaxy.univ-perp.fr/, Perpignan, France; Dugé de Bernonville et al. 2022) using *Populus trichocarpa* (v3.1) as a reference genome. Reads were first trimmed with *Trim Galore* (Galaxy v0.4.3.1) prior to mapping with *BSMAP* (Galaxy v1.0.0, Xi and Li, 2009) using default settings. Average sequencing depth after the mapping step ranged from 7X to 26X depending on genotypes (Table S1). Methylation calling was then achieved with *BSMAP methylation caller* (Galaxy v1.0.0) for the detection of methylated cytosines hereafter called SMPs for single methylated polymorphisms in the three methylation contexts (CpG, CHG and CHH). Bisulfite non-conversion rate ranked from 0.4 to 1.1% (Table S1). The *Methylkit* and *genomation* R packages (v1.18.0) were used for the analysis and annotation of DNA methylation data. SMPs were annotated for gene promoters (400 bp centered on the transcription start site (TSS) which allow to target the proximal promoter containing primary regulatory elements), genic regions (exons and introns), intergenic regions and transposable elements (TEs). With methylation expressed in percentage (%), we also used the rbd (read by density) normalization approach (Bellec et al. 2023) to study the link between DNA methylation and gene expression, with rbd = mCs x ratio (0-1) and where ratio = mCs / (mCs + Cs). MCs correspond to the number of methylated reads and Cs the number of un-methylated reads. Gene Ontology (GO) annotation was inferred from the *Arabidopsis* TAIR10 gene annotations using the best blastN hit (BlastN V3.0). GO terms Enrichment was performed using the *metascape* software with default parameters (Zhou et al. 2019).

### RNA-seq data recovery

The full set of RNA-seq data has already been published by Chateigner et al. 2020. We retrieved the same 20 genotypes analyzed here for WGS and WGBS and performed TPM normalization (Transcript Per Million, edgeR v3.26.4) for the comparison of different set of genes allowing to address gene size differences.

### Construction of phylogenetic and methyl-phylogenomic trees

Phylogenomic trees based on DNA methylation profiles were constructed with the *methylkit* software. Bed files corresponding to the methylation data for each of the genotypes were merged to form a methylation matrix grouping all genotypes by the context of methylation, tolerating up to 30% of missing data. For the CpG context only (symmetric case), the reads supporting the two strands were merged to improve coverage. Methylated matrices for each context were then filtered with SNPs data from the WGS to discard methylation calls due to genetic C/T polymorphisms. We set a minimum coverage of 7X for all genotypes and discarded the STR-010 genotype from the Rhin population as it did not reach this minimum value. The genotypes were clustered based on the similarity of their methylation profiles for each methylation context separately using ward hierarchical clustering and pearson’s correlation distance implemented in the *methylkit* software. Similarly, the phylogenetic tree was built using only SNPs without any missing value and with a minor allele frequency (MAF) above 5%. The genomic relationship matrix (GRM) was then estimated following VanRaden (2008) and the population structure evaluated by performing a hierarchical ascendant clustering using the Ward method on the GRM, converting relationships into dissimilarities.

### Detection of epigenomic signatures of local adaptation and/or drift

We took advantage of the available *pcadapt* tool (v4.3.3) in order to detect epigenomic markers involved in biological adaptation (Luu et al. 2017). We focused only on SMPs located in genic features (exons and introns) including promoter regions (200 bp around the TSS). Epigenetic markers (within genes) with particular patterns (i.e. outliers) were identified with the pool option of *pcadapt*. We divided the methylation matrices by 100 so that DNA methylation data are similar to frequencies. For all omics data we ran the analysis with the default options of *pcadapt* (MAF at 0.05). The number of principal components (PCs, K) captured varied between the three methylation contexts and was fixed according to the scree plots considering Catell’s rule (Cattel R.B, 1966). Thus, only principal components above the point of inflection were taken. For methylated CpGs, three components were retained while four components were selected for non-CpG contexts (CHG and CHH). We set the minimum cut-off for outlier detection at a p-value below 0.05 using Benjamini-Hochberg correction (FDR). Epigenetic markers that have disproportionately high contribution to the structure defining PCs were considered as putative markers of local adaptation. Using a different approach in order to characterize which epigenetic markers mimic the phylogenetic tree, we ran differential analysis between populations. First, we applied a filter on the standard deviation per position to select the cytosines which vary the most between the samples. Using the R package *MethylKit*, we fit a weighted fractional Logistic regression model to explain the ratio of methylated cytosines by the population structure. Weights are defined as the read coverage i.e., the sum of methylated and non-methylated cytosines after bisulfite conversion. The Chi-square Test is then used to assess the significance of the association between the cytosines and the genetic population structure. P-values are corrected by the Bonferroni approach and the significance threshold is fixed to 0.01. The comparison between the epigenetic trees built on top-markers and the phylogenetic tree are given by the tanglegram plots combining the two dendrograms for each context.

### Inference of duplicated genes and genomic fractions following polyploidization

The poplar genome was compared (with blastP) to the inferred Ancestral Eudicot Karyotypes (AEK) consisting of post-γ AEK with 21 proto-chromosomes and 9,022 ordered protogenes and a pre-γ AEK with 7 proto-chromosomes and 6,284 ordered protogenes, with γ being the shared triplication at the basis of rosids. The complete method for ancestral karyotype reconstruction is published in Murat et al. 2017. Filtering out one-to-two gene relationships between respectively AEK and poplar allowed the identification of duplicated genes inherited from the poplar-specific duplication within the *Salicaceae* family (Murat et al. 2015, Bellec et al. 2023). For each ancestral region, we detected compartments of the genome that underwent different fractionation in ancestral gene retention following polyploidization, defining the LF (Least Fractionated) and MF (Most Fractionated) compartments in the poplar genome, as proposed in Bellec et al. 2023. Differences between the evolutionary structural features (i.e., LF-vs. MF-genes) were assessed using 4 statistical tests: 2 distribution tests (Kolmogorov-Smirnov and Anderson-Darling tests), Kruskal-Wallis (median comparison) and T-test (mean comparison). Significant results were considered when at least two statistical tests passed the p-value cut-off (0.05). Differentially expressed genes (DEGs) and differentially methylated genes (DMGs) between the duplicated genes were investigated with the R package edgeR (v3.38.4, Chen et al. 2017) using TPM (to account gene size differences) and rbd (Bellec et al. 2023) data respectively. Differences were assessed using the likelihood ratio test and p-values adjusted by Benjamini-Hochberg method to control the false discovery rate.

## AUTHOR CONTRIBUTIONS

MDS and JS wrote the manuscript. All authors edited and helped to improve the manuscript; VB, CBuret, MCLD, ILJ and AD performed the sampling; DNA and RNA extraction were realized by VB, CBuret, MCLD, AD and MDS; Genomic data production was realized by JT, CD, CBesse; Bioinformatic and statistical analysis were done by JT, AG, OR, ASR, ILK, SM, EM, CA, VS, ES and MDS; SM and JS assumed the coordination; All authors read and approved the final manuscript.

## ACKNOWLEDGEMENTS

We are grateful to the to the genotoul bioinformatics platform Toulouse Occitanie (Bioinfo Genotoul, https://doi.org/10.15454/1.5572369328961167E12) and the Mesocentre Clermont Auvergne bioinformatics platform (https://doi.org/10.18145/aubi) for providing computing and storage resources and to the GBFOR, INRAE, 2018, Forest Genetics and Biomass Facility (https://doi.org/10.15454/1.5572308287502317E12) for the experimental design setup and samples collection. We thank COST action (European Cooperation in Science and Technology) EPIgenetic mechanisms of Crop Adaptation To Climate cHange (EPICATCH; grant number CA19125) for active discussion.

## DATA AVAILABILITY STATEMENT

The raw data for WGS, WGBS and RNAseq are stored in the NCBI website under the following accession numbers PRJNA818172 BioProject (WGS), PRJNA828400 BioProject (WGBS), GSE128482 (RNA-seq). The processed SNPs, methylation matrices, LF/MF, duplicated genes, pcadapt, hypo/up and Hyper/Down candidate genes can be found at: https://entrepot.recherche.data.gouv.fr/privateurl.xhtml?token=0d8bacbb-71aa-4352-9da4-cc6ed16a6190.

## SUPPORTING INFORMATION

**<u>Figure S1.</u>** Global DNA methylation percentage distribution between the 20 *P. nigra* genotypes in CpG, CHG and CHH contexts.

**<u>Figure S2.</u>** DNA Methylation coverage on genes and TEs in the CpG, CHG and CHH contexts.

**<u>Figure S3.</u>** Gene ontology annotation using biological process terms of LF, MF and duplicated genes.

**<u>Figure S4.</u>** Venn diagram of differentially expressed genes (DEGs) and differentially methylated genes (DMGs) between the duplicated genes.

**<u>Figure S5.</u>** Phylo-epigenomic tree reconstruction for DNA methylation in CHG and CHH contexts.

**<u>Figure S6.</u>** Methylation markers involved in local adaptation.

**<u>Figure S7.</u>** DNA methylation dynamics of disease resistance (R) genes in the 20 black poplar trees.

**<u>Figure S8.</u>** Comparison between gene expression and DNA methylation.

**<u>Table S1.</u>** Mapping and methylation statistics for 20 black *P. nigra* genotypes.

